# Mind blanking is a distinct mental state linked to a recurrent brain profile of globally positive connectivity during ongoing mentation

**DOI:** 10.1101/2021.05.10.443428

**Authors:** Sepehr Mortaheb, Laurens Van Calster, Federico Raimondo, Manousos A. Klados, Paradeisios Alexandros Boulakis, Kleio Georgoula, Steve Majerus, Dimitri Van De Ville, Athena Demertzi

## Abstract

Mind blanking (MB) is a waking state during which we do not report any mental content, challenging the view of a constantly thought-oriented brain. Here, we comprehensively characterize the MB’s neurobehavioral profile with the aim to delineate its role during ongoing mentation. Using fMRI experience-sampling, we show that MB is reported less frequently, faster, and with low transitional dynamics among other mental states, pointing to its role as a transient mental relay. Regarding its neural underpinnings, we observe higher global signal amplitude during MB reports, indicating a distinct physiological substrate. Using the time-varying functional connectome MB reports get classified with high accuracy, suggesting that MB has a unique neural composition. Indeed, a pattern of globally positive-phase coherence shows the highest similarity to the connectivity patterns associated with MB reports. We observe that this pattern’s rigid signal architecture hinders content reportability due to the brain’s inability to differentiate signals in an informative way. Collectively, we show that MB has a unique neurobehavioural profile, indicating that non-reportable mental events can happen during wakefulness. Our results add to the characterization of spontaneous mentation and pave the way for more mechanistic investigations of MB’s phenomenology.

**Significance Statement:** The human mind is generally assumed to be thought-oriented. Mind blanking (MB) challenges this stance because it appears as if we are derived of any particular mental content to report. We here show that, during spontaneous thinking, MB is a mental state that happens by default, it has a unique behavioural profile, and it is of a rigid neural architecture that does not permit the formulation of reportable contents. Our work essentially proposes that non-reportable mental events can happen during wakefulness, and challenges the view of the mind as a constant thought-oriented operator.

## Introduction

During ongoing task-free conditions, spontaneous experience is ongoing, dynamic, and rich in mental content (1), taking the form of mental states. Mental states are transient cognitive or emotional occurrences that are described in terms of particular content (what the state is ‘about’) and the relation we bear to this content (e.g. imagining, remembering, fearing). Thoughts, in that sense, are sequences of mental states (2). Ongoing experience can also show moments of mind blanking (MB) during which there is a failure to report the content of thoughts, often accompanied by a post-hoc realization that our mind “went away” (3). Behavioral studies indicate that MB happens scarcely during task performance, yet with a considerable frequency. For example, it has been shown that, during focused tasks, MB was reported on average 14.5%of the times whenever subjects evaluated their mental state upon request (3), and 18% of the time when participants reported MB by self-catching (4). In terms of neural correlates, instructing participants to “think of nothing” as compared to “let your mind wander” led to lower fMRI functional connectivity between the default mode network (DMN) and frontal, visual, and salience networks. On the one hand, this is indicative of how pre-scan instructions influence connectivity results (5), and how MB might exhibit a distinct neural profile on the other. Indeed, MB has also been associated with deactivation of Broca’s area and parts of the hippocampus, as well as with activation of the anterior cingulate cortex, which has been interpreted as evidence for reduced inner speech (6). Decreased functional connectivity in the posterior regions of the DMN and increased connectivity in the dorsal attentional network has also been found in an experienced meditator practicing content-minimized awareness, which can be considered as a phenomenological proxy to MB (7).

Collectively, these studies indicate that, although the investigation of MB is rising over the years, its neurobehavioral characterization remains inconclusive. This might be due to several reasons. First, MB has been studied after deliberately inducing it or in highly-trained individuals, therefore its spontaneous occurrences are not generalisable. Second, in some cases MB has been studied in isolation from other mental states, therefore its inter-state dynamics are lacking. Third, current MB’s neural correlates concern a limited number of brain regions, leaving the whole-brain functional connectome uncharted. In this study, we aimed at delineating the neurofunctional profile of MB in a comprehensive way. For this purpose, we used fMRI-based experience-sampling in typical individuals (8) in order to: a) account for the behavioral quantification of spontaneous (no induction) MB occurrences, b) determine MB’s inter-mental-state dynamics, and c) estimate the MB’s functional fine-grained connectome at the whole-brain level.

## Results

We used previously acquired data (8) collected from 36 healthy participants (27 females, 9 males, mean age: 23y±2.9) within a 3T MRI scanner while they were at rest with eyes open. The experience-sampling task concerned randomly presented auditory sounds (n=50) that prompted the participants to evaluate and chose by button press the mental content in which they were prior the probe. Possible mental states were: Absence (i.e., MB), perception of sensory stimuli (Sens), stimulus-dependent thoughts (SDep), and stimulus-independent thoughts (SInd) (Fig 1).

**Figure 1.**
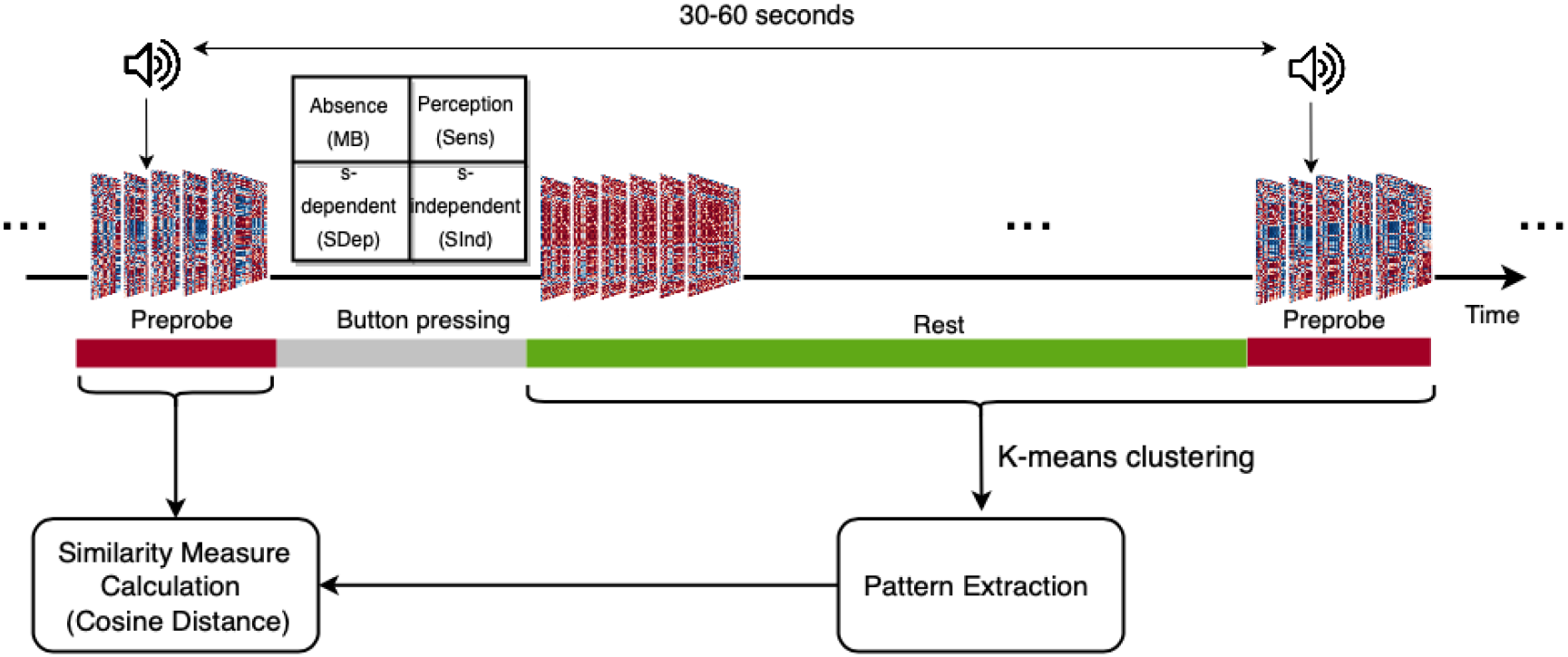
Data acquisition and analysis paradigm. While at rest, participants were interrupted by an auditory probe to report their immediate mental state by choosing with button press among: i) Absence (MB), ii) Perception (sensory perception), iii) stimulus-dependent thoughts (thoughts related to the immediate environment), and iv) stimulus-independent thoughts (thoughts unrelated to the immediate environment). By means of Hilbert transform and phase-based coherence analysis, the connectivity matrices at each volume were estimated. Considering the effect of hemodynamic response, 2 pre-probe and 3 post-probe connectivity matrices were considered as the analysis window. The matrices were then concatenated across all subjects. By means of k-means clustering, they were summarized into recurrent brain patterns. The cosine distance was used as a similarity measure between the report-related connectivity matrices and the brain patterns.

### Behavioral analysis

Considering the occurrence rate over time, MB was reported significantly fewer times than the other mental state (median=2.5, IQR=3, min=0, max=9; Fig. 2A). With respect to reaction time, there was a main effect of mental states *χ*^2^(3)=66.63, p<0.001; generalized linear mixed model analysis; Fig. 2B), with MB being reported faster than Sdep (z=3.81, p=0.0008) and SInd (z=3.37, p=0.0042) but with no significant differences from Sens (z=-0.73, p=0.89; post-hoc Tukey test). The evaluation of the dynamic transitions among different mental states showed exceptionally low but equal probabilities (0.06) for reporting MB when departing from a content-oriented state (Fig. 2C). Also, the probability of re-reporting MB was particularly low (0.04). Finally, the hypothesis of a uniform distribution of reports across the session could not be rejected either for MB *χ*^2^(9)=12.31, p=0.20, *φ*=0.35) or SDep *χ*^2^(9)=5.25, p=0.81, *φ*=0.10) or SInd *χ*^2^(9)=4.22, p=0.90, *φ*=0.07). Sens reports, though, were not uniformly distributed over time (*χ*^2^(9)=18.15, p=0.03, *φ*=0.23; Fig. S1).

**Figure 2.**
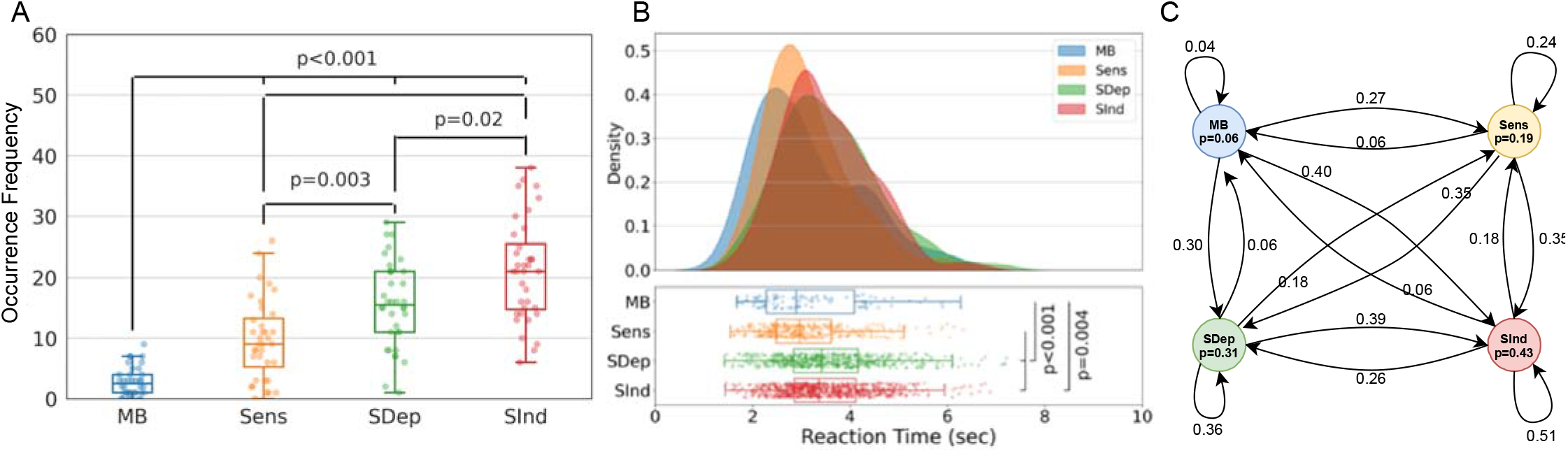
MB is characterized by a distinct behavioral profile. A) MB shows significantly low reportability by comparisonto the other mental states (FDR p<0.05). B) MB is reported significantly faster than SDep and SInd mental states. Yet, thereaction time were not significant comparing MB and Sens reports. Due to the tail of the reaction time distributions, ageneralized mixed model with a gamma distribution was fitted to the data for statistical analysis. P-values are from the post-hocTukey test performed after finding significant effect of mental states on the reaction times. C) A Markov model shows thatreporting a MB state, after any other mental states is low but equal (6%), suggesting that MB might serve as a transient mentalrelay during spontaneous mentation.

### Functional MRI analysis

#### MB is supported by a distinct physiological state

In order to estimate the MB’s functional connectome, we first sought to delineate the contribution of the global signal (GS). This was because the GS has been previously shown to contain neural sources (9–11) and, thus, can be of functional significance. The spatially averaged timeseries were extracted from the connectome’s ROIs and its amplitude was estimated within a total of five volumes per probe: the two volumes preceding the probe and three volumes after it (Fig.1). The selection of this time window was justified by the repetition time (TR, 2sec) and the hemodynamic response which reaches its maximum after 3 scans post-event (12). This analysis showed a significant effect of mental state on the GS amplitude (*χ*^2^(3)=12.474, p=0.006; generalized linear mixed model analysis), with higher amplitude relating to the volumes surrounding MB reports as compared to those linked to SDep (z=3.3, p=0.005) and SInd reports (z=2.55, p=0.05; post-hoc Tukey test; Fig. 3). Similar results were obtained when the analysis window lagged between zero frames (i.e., 5 scans preprobe) up to 3 volumes (i.e., 2 pre-probe, and 3 post-probe scans; Fig. S2). As the GS contributes deferentially to the reportability of mental states, we decided to include it in the connectivity analyses. For comprehensive purposes, all subsequent analyses were also performed without the GS as well (see below).

**Figure 3.**
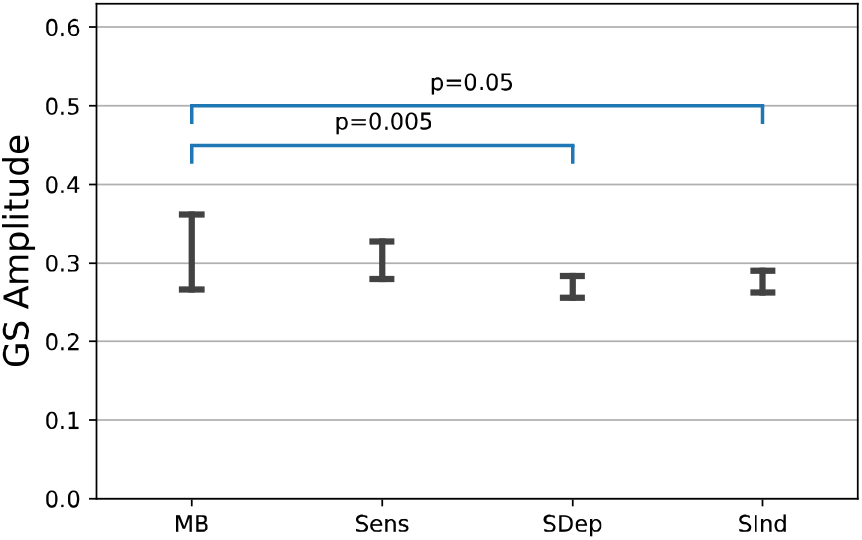
MB-labelled volumes are characterized by high global signal amplitude. The average absolute value of theglobal signal estimated on 2 pre-probe and 3 post-probe scans shows that the GS amplitude is significantly higher for thosevolumes reporting MB compared to the amplitude of the global signal observed in volumes reporting content-oriented states.Notes: Bars show the mean absolute value and error bars show 95% confidence interval.

#### MB is accurately classified by means of phase-based coherence

To check whether MB is of distinct neural profile, we first tested whether it can be classified from other mental states using the functional connectome. The Hilbert transform was used to estimate frame-wise phase-based coherence matrices within the above-mentioned 5-volume window. Considering these connectivity matrices as feature vectors (5 vectors per probe), a support vector machine (SVM) classifier with a 5-fold cross-validation with 10 repeats classified MB reports from all mental states with an average precision of 1, average recall of 0.79, and average balanced accuracy of 0.89. In addition, a one-vs-one strategy to classify MB from the other reports separately led to high classification performance (Table 1). To compare the results with an empirical chance level, a dummy classifier was further used to separate MB-labeled matrices from the matrices corresponding to the other mental states. This dummy classifier generated random predictions by respecting the training set class distribution. Collectively, by comparing all the performance metrics of the MB classification using SVM and the dummy classifier, we found that the SVM successfully separated the functional connectomes of MB reports from those belonging to the other mental states.

**Table 1.** Performance of SVM classifier when predicting MB reports based on phase-based coherence matrices. CI shows the0.95% confidence interval

|  | Balanced Accuracy | Recall | Precision |
| --- | --- | --- | --- |
| ***MB vs. Sens*** | 0.97, CI=(0.87, 1.07) | 0.94, CI=(0.73, 1.15) | 0.99, CI=(0.97, 1.03) |
| ***MB vs. SDep*** | 0.95, CI=(0.83, 1.07) | 0.91, CI=(0.67, 1.14) | 1, CI=(1,1) |
| ***MB vs. SInd*** | 0.94, CI=(0.79, 1.09) | 0.87, CI=(0.58, 1.17) | 1, CI=(1,1) |
| ***MB vs. Others*** | 0.90, CI=(0.73, 1.07) | 0.79, CI=(0.45, 1.13) | 1, CI=(1,1) |
| ***MB vs. Others (dummy)*** | 0.50, CI=(0.43, 0.57) | 0.05, CI=(-0.07, 0.18) | 0.06, CI=(-0.10, 0.22) |

#### Functional connectomes organize into distinct recurrent brain patterns

Under the hypothesis that the MB’s neural signature is contained in connectivity dynamics, we investigated how the frame-wise functional connectome organizes into distinct connectivity patterns. By concatenating all the estimated connectivity matrices across subjects and by applying k-means clustering, we determined four main functional brain patterns which appeared recurrently across the resting state periods, replicating previous results (13) in spite of different acquisitions parameters and parcellation schemes. The patterns were characterized by distinct signal configurations: a pattern of complex inter-areal interactions, containing positive and negative phase coherence values between long-range and short-range regions (Pattern 1), a pattern showing anti-correlations primarily between the visual network and the other networks (Pattern 2), a pattern with overall positive inter-areal phase coherency (Pattern 3), and a pattern of overall low inter-areal coherency (Pattern 4; Fig. 4A). In terms of occurrence rate, Pattern 4 appeared at a significantly higher rate than Pattern 1 (t(35) =7.131, p<0.001, Cohen’s d=1.18), Pattern 2 (t(35) =7.495, p<0.001, Cohen’s d=1.25) and Pattern 3 (t(35) =5.857, p<0.001, Cohen’s d=0.98, p-values are FDR corrected at α=0.05; Fig. 4A). These patterns also emerged when using different cluster size (ranging from 3 to 7) and different analysis window lags (ranging from zero up to 3 frames). The significantly high occurrence rate of the low inter-areal connectivity pattern was observed across all cluster sizes and window lag combinations (Fig. S3-S6).

**Figure 4.**
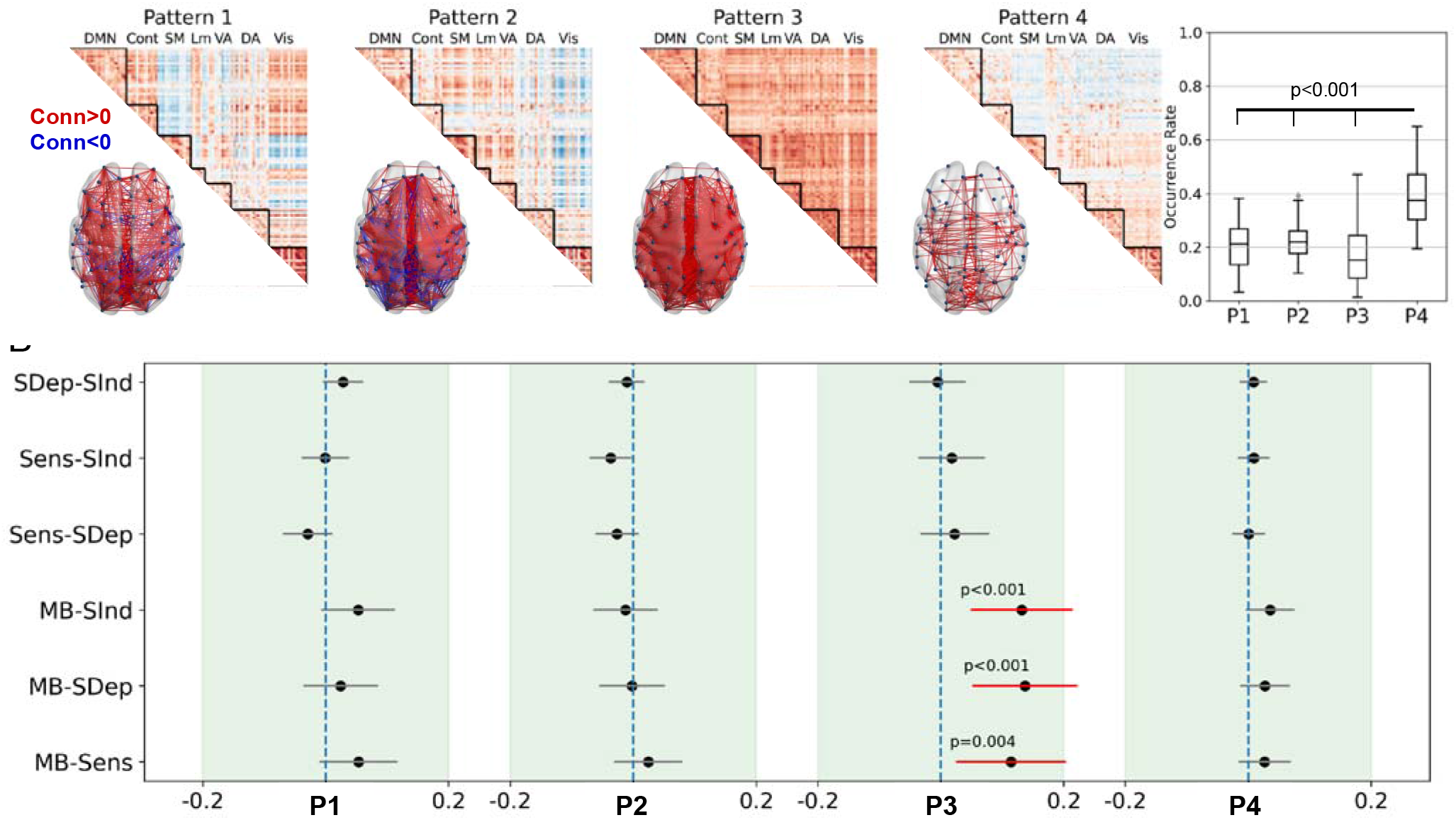
Neurobehavioral coupling analysis shows that MB is associated with an overall positive inter-regional brainconnectivity pattern. A) Brain functional organization during rest can be summarized into four main connectivity patterns of complex cortical interaction (Pattern 1), visual network anticorrelation (Pattern 2), globally positive coherency (pattern 3), and low interareal connectivity (pattern4), with highest occurrence rate related to the fourth pattern. B) The globally positive phase-coherence Pattern 3 shows the highest similarity to the connectivity matrices related to MB reports compared to the other mental states. Black dots show the contrast between similarity measures of related mental states; error bars indicate 95% confidence interval. A generalized linear mixed model (gamma distribution, log link function, mental states as fixed effect, andsubjects as random effect) was fitted to the similarity measures of each pattern separately. In case of significant effect of themental states after Bonferroni correction for multiple tests 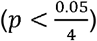, a Tukey post-hoc test was performed to test pairwise mental state differences.

### Neurobehavioral coupling

To determine which brain pattern was the closest to the MB reports, we used the cosine distance as the similarity measure between five connectivity matrices prior to each report (i.e., analysis window) and the four resting brain patterns (Fig. 1). Using a generalized linear mixed model fit to the distance measures of each brain pattern separately, we found a significant effect of mental state for distance values to Pattern *χ*^2^(3) =19.088, p=0.0002). Pattern 3 further showed higher similarity to MB compared to the reports of Sens (Estimate=0.114, low CI=0.027, high CI=0.202, p=0.004), SDep thoughts (Estimate=0.137, low CI=0.053, high CI=0.221, p=0.0002), and SInd thoughts (Estimate=0.132, low CI=0.050, high CI=0.213, p=0.0002; Post-hoc tukey tests; Fig 4B). These results were also replicated using different analysis window lags (Fig. S16-S19).

For comprehensive purposes, a supplementary analysis of the neurobehavioral coupling was performed by omitting the GS by subtraction and by regression. Global signal subtraction (GSS) refers to subtracting the GS from the ROI preprocessed timeseries. Global signal regression (GSR) concerns the removal of the average functional connectivity values from the preprocessed ROI timeseries via linear regression. After applying GSS and GSR, all brain patterns were reproduced, except for Pattern 3 (Fig. S15A). The same observation was noticed on the clustering results with different cluster size (Fig. S7-S14). The overall effect of GSS and GSR on the connectivity patterns was the shift of connectivity value distributions towards negative values (Fig. S15B) and this shift was more prominent for Pattern 3. In addition, inter-pattern correlation analysis showed that Pattern 3 had the lowest similarity to itself after GSS and GSR (ρ=0.59; Fig. S15C). Considering a 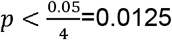 threshold to correct for multiple tests, no significant effect of mental states on the similarity measures were found for any pattern (GSS(Pattern 1: p=0.931, pattern 2: p=0.116, pattern 3: p=0.294, pattern 4: p=0.573), GSR(Pattern 1: p=0.109, pattern 2: p=0.022, pattern 3: p=0.276, pattern 4: p=0.093); Fig. S20 and S24). These results suggest that the GS carries partially independent neural information and contributes to the cerebral profile of MB reportability.

## Discussion

We used experience-sampling paired with fMRI to determine the neurobehavioral profile of MB in typical individuals. Collectively, our results show that MB is a unique mental state supported by a distinct neural architecture which contributes meaningfully to spontaneous mental activity.

Behaviorally, we found that individuals report fewer MB occurrences that are reported faster than other mental states. This finding is in line with previous studies showing that MB gets reported significantly less often than thought-related states (3, 4), although the opposite effect was also reported (6). These discrepancies might be attributed to the study protocol, where in the latter study participants were encouraged to stay engaged into thinking about nothing (6). This implies that MB might be a flexible and trainable mental state, which, once introduced as an option, it can be informative of one’s ongoing phenomenology. Our results also align with studies reporting similarly fast MB reaction times while participants are involved in sustained attention to response tasks (14, 15). Other investigations, though, show that MB can be reported more slowly compared to other mental states, which was interpreted as MB facilitating sluggishness in responses or as the result of decreases in alertness and arousal during task performance (17). Here, we consider that the fast reaction times for MB and the longer reaction times for thought-oriented mental states (Sdep, Sind) might be attributed to an additional cognitive evaluation of the latter. In other words, when thoughts are occupied by specific content, this is translated in longer cognitive evaluation as to the particularities of this content.

This stance implies that MB can be a mental state that is “content-free” and, as such, it is reported faster. This interpretation is supported by previous investigations using self-paced focused reading with self-catches of MB and mind wandering (3). Although this is a tempting consideration, we recognize that the content-free nature of MB reports could not be directly addressed here. Attempting to delineate the mechanisms of MB, we indeed observe that attention can act as a mediating process that drives content reportability (18), so that participants do entertain content-full thoughts but fail to attend to them, therefore leading to attentional lapses (15, 17, 19). At the same time, it can be that MB is a matter of participants’ meta-cognitive capacities, in that MB is more about a “cognitive-evaluation free” or “meta-awareness free” mental state rather than lack of mental content. Equally, MB might not be a state of no experience. Rather, it can work as a “transition mode” between modifications of experience (content), as we move from one state to the other. This scenario suits our results of the low probabilities of MB to get reported when previously in another mental state. In that case, departures from MB are more likely to lead towards thought-oriented reports, and less likely to return to MB. However, these findings should be considered within the temporal constraints of the experience-sampling paradigm. This means that one cannot assume that this dynamic sequencing reflects actual mental state transitions because the temporal structure between the reports is not continuous. Consequently, one cannot ensure whether other mental states appeared between reports. Despite this limitation, the finding that the equally small probabilities to report MB when in another state and vice-versa indicates that MB might not be driven by any specific mental content, therefore serving as a transient mental relay (20). This means that thoughts with reportable content can lead towards more mental contents due to semantic associations, hence creating the perception of a stream of consciousness (2). Since MB is not semantically associated with any particular mental content, it may therefore occur scarcely during ongoing experience. As such, phenomenologically “empty” mental states might have less of an anchoring effect than content-full states. Finally, our finding of a uniform distribution of MB reports over time, also reported elsewhere (3, 21), further suggests that MB happens spontaneously across time and is not an artifact of fatigue which would lead to more occurrences at the end of the recordings. Taken together, the behavioral results support that MB is a distinct mental state with a unique position among thought-oriented reports. In order to shed light on the refined mechanisms underlying MB reportability we further suggest that future work addresses MB in terms of content, attention, and metacognitive capacities.

In terms of MB’s neural underpinning, we show four distinct brain patterns which re-occur dynamically during the resting periods of the experience-sampling task. These brain patterns bear great resemblance with what we previously reported as recurrent brain configurations during pure resting state fMRI acquisitions across healthy individuals and brain-injured patients (13). The fact that these patterns appear across independent datasets, also in non-human primates (22), utilizing different paradigms, different brain parcellations, and different cluster sizes points to their universality and robustness. The finding that the pattern with an overall low inter-areal connectivity (Pattern 4) shows the highest occurrence probability in comparison to the other patterns, can be explained by the fact that this configuration has the highest similarity to the underlying structural connectome (13). As such, this pattern may act as a foundation upon which the others can occur, by showing divergence of function from structure, which is linked to mental flexibility (23).

At the same time, the pattern with the all-to-all positive inter-areal connectivity (Pattern 3) had the lowest occurrence probability and the highest similarity to the connectivity matrices preceding MB reports. Such high prevalence of comparable signal configurations was previously shown during NREM slow-wave sleep, wherein overall minimal neuronal firing was translated as globally positive connectivity (24, 25). Studies in rats (26) show that such periods of neuronal silencing can happen also during wakefulness in the form of neuronal firing rate reduction leading to slow wave activity, which is indicative of local sleeps. When applied to humans, it has been argued that these instances of local sleeps can be the phenomenological counterpart of MB (16). In that respect, wakefulness does not only support constantly on-periods of neuronal function. Rather, our brains can also show instances of neural down-states even during wakefulness, possibly for homeostatic reasonse (27), which can be translated as global positive connectivity and phenomenologically interpreted as MB.

Importantly, the Pattern’s 3 architecture demolished towards negative coherence values once the global signal (GS) was regressed out. Here, the dramatic effect of GS removal on the dynamic functional connectome, also previously reported (28), was expected as it is known that this process enhances functional anticorrelations by shifting the distribution of correlation values in the negative direction (29–31). In that respect, removal of the GS is a debatable issue for resting state analyses (32), whether to keep it or regress it out. To date, there is support for both views. The GS has been shown to have a neuronal counterpart (28, 33) which promotes behavior (9). And it is shown to reflect fMRI nuisance sources such as motion, scanner artifacts, respiration (34), cardiac rate (35) and vascular activity (36, 37). In our analysis we eventually included the GS because we found that its amplitude preferentially dominated the scanning volumes associated to MB reports. The supplementary analysis of the neurobehavioral coupling without the GS confirmed that the GS contributes meaningfully to the phenomenology of MB as it dramatically demolished the overall inter-regional positive coherence of Pattern 3. At the moment, we can only speculate about what the high GS amplitude might mean for MB reportability. In terms of physiological relevance, spontaneous GS amplitude was found to correlate negatively with EEG vigilance (alpha, beta oscillations), while increases in EEG vigilance due to caffeine ingestion were associated with reduced GS amplitude (38). In macaques, electrocorticography showed that widespread transient and synchronous cortical activity was linked to low arousal in a series of sequential spectral transitions, from a decrease in mid-frequency activity, accompanied by increase in gamma band, to be eventually followed by increases in delta band (39). When these transient events in animals were linked to fMRI motifs in humans, there was a close association between the GS and these events as well as with a subcortical involvement in both features, which was considered as consistent with an origin in arousal (40). These results, jointly with the here elevated GS amplitude during MB, further support the possibility of neuronal silencing during wakefulness as discussed above. In the absence of a mechanistic investigation, though, this remains an ongoing hypothesis. We nevertheless wish to suggest that future connectivity analyses are performed both with GS included and removed in order to account for its unexpected effects.

Theoretically-wise, how can MB reports happen? To date, it seems that MB further challenges the boundaries of various models of conscious experience. For example, the Global Neuronal Workspace Theory (GNWT) (41) posits that a stimulus becomes reportable when some of its locally processed information becomes available to a wide range of brain regions, forming a balanced distributed network (42). A key process of this global broadcasting is ignition (43). Ignition is characterized by the sudden, coherent, and exclusive activation of a subset of workspace neurons which code a particular content, while the remainder of the workspace neurons stay inhibited. If the GNW ignition is always related to selective neural activation and inhibition (content), the theory cannot account for how MB can still be reported if it is linked to a functional connectome with only positive connections. This is similar for the Integrated Information Theory (IIT) (44). According to it, in order to generate an experience a physical system must be able to discriminate among a large repertoire of states (i.e., information). This must be done as a single system that cannot be decomposed into a collection of causally independent parts (i.e., integration). So far, the IIT embraces the inability to report mental content in brain states with extreme functional integration, like during generalized epilepsy (45). In such a brain state, an abnormally large number of regions work in synchrony, and, as a result, the brain becomes no longer capable of processing information in a way that leads to conscious experience. The here identified all-to-all positive connectivity pattern shows the highest level of integration and efficiency and the lowest level of segregation and modularity compared to the other brain patterns (13). Therefore, this may imply that such a neural configuration is unable to produce a balance between values of integrated information and segregation of it, leading to limited experiences, such as MB. If the role of integration is emphasised over the role of segregation, such as in the recent version of IIT, then MB challenges that approach, making a clear case for the importance of segregation of information within neural configurations of conscious content. Importantly, though, the integration in IIT happens only when there is a content of experience, being reported or not, which is totally counterintuitive for MB. Both theories essentially start from the premise that experience is made up of various bits from which a unified experience arises. As MB does not provide such building blocks, it seems to be a kind of global state of unified experience, and conscious content being the modifications of such a basal conscious field, according to Searle’s unified field model (46). If this interpretation is considered, then the current findings pose important challenge to building block models of conscious experience.

Our analysis leaves several questions unaddressed. First, the current design does not permit to determine the underlying mechanism that drives MB, i.e., whether it is an effect of attention, memory, or language. Such determination is expected to shed light on MB’s modulatory mechanisms as well, and therefore further decipher its functional significance in variant conditions. Second, apart from the intrinsic problems with the validity and reliability of self-reports during experience-sampling (47), we also utilized a probe-catching methodology. This means that participants were interrupted during spontaneous thinking by a probe, asking them to choose an appropriate option to describe their thought-state. Such a probe-framing technique can restrict the estimation of potential phenomenological switches happening in between. Indeed, as the probes were appearing in pre-determined time points we cannot exclude the possibility of mental contents happening during the inter-probe intervals, and hence they were missed to be reported. Also, probe-framing can be suboptimal in capturing spontaneous thinking because it might lead to an inflated number of MB reports. This is because participants could choose this category since it was pre-established, which they could otherwise not report if they were to identify spontaneously (48). However, given that MB occurrences were not reported with a comparable high frequency to the content-oriented states might indicate that MB was evaluated in a representative way across the evaluation, leading to infrequent occurrences across participants. Finally, the high TR during the fMRI acquisition (2.04s) could also echo the temporal implications of the MB profiling. By means of simultaneous EEG-fMRI recordings, more light is expected to be shed on fine-grained temporal dynamics of MB. Such simultaneous multi-modal recordings are expected to also illuminate the assumption of slow-wave activity as the corresponding neural mechanism of MB.

In conclusion, our study suggests that MB can be considered as a default mental state occupying a unique position among thought-oriented reports. Its rigid neurofunctional profile could account for the inability to report mental content due to the brain’s inability to differentiate signals in an informative way. While waiting for the underlying mechanisms of MB to be illuminated, these data suggest that instantaneous non-reportable mental events can happen during wakefulness, setting MB as a prominent mental state of the phenomenology of ongoing experience.

## Materials and Methods

### Dataset

Thirty-six healthy right-handed adults participated in an fMRI experience-sampling task (8). This sample size has been shown sufficiently reliable for group-level fMRI (49). All participants gave their written informed consent to take part in the experiment. Ethics committee of the University Hospital of Liège approved the study. Data were acquired during resting state while participants were lying inside the scanner with eyes open. At random times, they were interrupted by an auditory tone, probing them to report their immediate mental state via button presses (Fig.1, Upper panel). The sampling probes were randomly distributed between 30 and 60 seconds. Each probe started with the appearance of an exclamation mark lasting for 1000 ms inviting the participants to review and characterize the cognitive event(s) they just experienced. Then, on the screen four categories for a broad characterization of the cognitive experiences were shown: Absence, Perception, Stimulus-dependent thought, and Stimulus-independent thought. Absence was defined as mind blanking or empty state of mind. Perceptions represented the acknowledgment of a stimulus through one or more senses without any internal thought. Thoughts were distinguished as stimulus-dependent (i.e. with awareness of the immediate environment), or stimulus-independent (i.e. with no awareness of the immediate environment). For reporting, participants used two response boxes, one in each hand. Participants used an egocentric mental projection of their fingers onto the screen so that each finger corresponded to a specific mental category. Depending on the probes’ trigger times and participants’ reaction times, the duration of the recording session was variable (48-58 min). To minimize misclassification rates, participants had a training session outside of the scanner at least 24 hours before the actual session.

### Imaging setup

Experiments were carried out on a 3-T head-only scanner (Magnetom Allegra, Siemens Medical Solutions, Erlangen, Germany) operated with the standard transmit–receive quadrature head coil. fMRI data were acquired using a T2*-weighted gradient-echo EPI sequence with the following parameters: repetition time (TR) = 2040 msec, echo time (TE) = 30 msec, field of view (FOV) = 192×192 mm^2^, 64×64 matrix, 34 axial slices with 3 mm thickness and 25% interslice gap to cover most of the brain. A high-resolution T1-weighted MP-RAGE image was acquired for anatomical reference (TR = 1960 msec, TE = 4.4 msec, inversion time = 1100 msec, FOV = 230×173 mm, matrix size = 256×192×176, voxel size = 0.9×0.9×0.9 mm). The participant’s head was restrained using a vacuum cushion to minimize head movement. Stimuli were displayed on a screen positioned at the rear of the scanner, which the participant could comfortably see using a head coil mounted mirror.

### Behavioral analysis

Analyses were performed using locally developed codes in Python and R. Six paired t-tests were used to compare the number of reports of each mental state across participants (p-values were FDR-corrected with a significance level of α=0.05). A generalized linear mixed model with a gamma distribution and inverse link function tested the relationship between reaction times and mental states (The choice of the generalized linear mixed model was because of positive tail in the distribution of reaction times and inhomogeneity of its variance across mental states). Mental state reports were considered as fixed effects and participants were considered as the random effects with sex and age as confound variables. In case of significant main effects, post-hoc Tukey pairwise comparisons were applied. To model dynamic transition between mental state reports, a Markov model was used to calculate the transition probabilities between participants’ reports over the experiment. The uniformity of the distribution of each report over the acquisition duration was tested using the *χ*^2^ test on the time point of reports across all participants. The acquisition duration of each subject was divided into 10 equal temporal bins and the number of reports at each bin was counted. To calculate the effect size of the *χ*^2^ test, *φ* measure was used (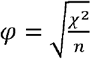, where n is the number of observations).

### fMRI preprocessing

Preprocessing and denoising were performed using a locally developed pipeline written in Python (nipype package (50)) encompassing toolboxes from Statistical Parametric Mapping 12 (51), FSL 6.0 (52), AFNI (53), and ART (http://web.mit.edu/swg/software.htm). Using this pipeline, all the functional volumes were realigned to the first volume and then, in a second pass, to their average. Estimated motion parameters were then used for artifact detection. An image was defined as an outlier or artifact image if the head displacement in the x, y, or z direction was greater than 3 mm from the previous frame, if the rotational displacement was greater than 0.05 rad from the previous frame, or if the global mean intensity in the image was greater than 3 SDs from the mean image intensity for the entire scans. After skull-stripping of structural data (using FSL BET (54) with fractional intensity of 0.3), realigned functional images were registered to the bias-corrected structural image in the subject space (rigid-body transformation with normalized mutual information cost function). After extracting white matter (WM), grey matter (GM), and cerebrospinal fluid (CSF) masks, all the data and masks were transformed into the standard stereotaxic Montreal Neurological Institute (MNI) space (MNI152 with 2 mm resolution). WM and CSF masks were further eroded by one voxel. For noise reduction, we modeled the influence of noise as a voxel specific linear combination of multiple empirically estimated noise sources by deriving the first five principal components from WM and CSF masked functional data separately. These nuisance regressors together with detected outlier volumes, motion parameters and their first-order derivative were used to create a design matrix in the first-level general linear model (GLM). After smoothing the functional data using a Gaussian kernel of 6-mm full width at half-maximum, the designed GLM was fitted to the data. Before applying GLM, functional data were demeaned and detrended and all the motion-related and tissue-based regressors were first normalized and then demeaned and detrended using the approach explained in (55). A temporal causal bandpass filter of 0.01 to 0.04 Hz was then applied on the residuals of the model to extract low frequency fluctuations of the BOLD signal. Schaefer atlas (56) with 100 ROIs were then used to parcellate each individual brain. Average of voxel time series in each region was considered as the extracted ROI time series and were used for further analysis.

### Functional connectivity patterns estimation

We used the phase-based coherence analysis to extract between-region connectivity patterns at each time point of the scanning session (13). For each participant *i*, after z-normalization of time series at each region *r* (i.e., *x_i,r_*(*t*)), the instantaneous phase of each time series were calculated using Hilbert transform as:

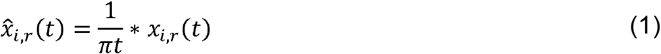

in which * indicates a convolution operator. Using this transformation, an analytical signal was produced for each regional timeseries as:

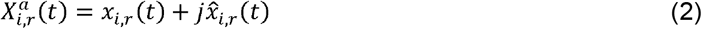

where 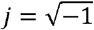. From this analytical signal, the instantaneous phase of each time series can be estimated as:

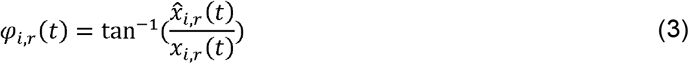

After wrapping each instantaneous phase signal of *φ_i,r_*(*t*) to the [-*π, π*] interval and naming the obtained signal as *θ_i,r_*(*t*), a connectivity measure for each pair of regions was calculated as the cosine of their phase difference. For example, theconnectivity measure between regions *r* and *s* in subject *i* was defined as:

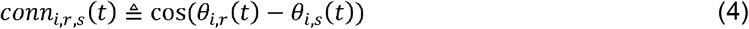

By this definition, completely synchronized time series lead to have a connectivity value of 1, completely desynchronized timeseries produce a connectivity value of zero, and anti-correlated time series produce a connectivity measure of −1. Using this approach, a connectivity matrix of 100×100 was created at each time point *t* for each subject *i* that we called it *C_i_*(*t*):

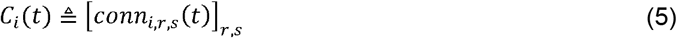

After collecting connectivity matrices of all time points of all participants, k-means clustering was applied on all the estimated connectivity matrices. With this technique, four robust and reproducible patterns were extracted as the centroids of the clusters and each resting connectivity matrix was assigned to one of the extracted patterns (We chose to extract four patterns to compare our results with our previous research (13). However, we replicated all the analysis using different number of clusters ranging from 3 to 7). The occurrence rate of each pattern was simply calculated by counting the number of matrices which were assigned to each specific pattern at each subject separately. Significant differences between patterns occurrence rates were analyzed using paired t-test and FDR correction of p-values over six possible pairwise comparisons.

### Classification of mind blanking based on time-varying connectivity matrices

Phase-based coherency matrices within the analysis windows were considered as the feature vectors and the related mental state reports as the class labels. First, a support vector machine (SVM) model for binary classification were designed to classify MB reports from all the other reports. As the dataset was imbalanced, we calculated precision 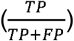, recall 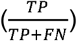, and balanced accuracy 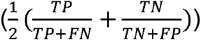 as the efficiency parameters of the classifier (TP: True Positives, TN: True Negatives, FP: False Positives, FN: False Negatives; MB reports were defined as positive class). As the cross validation strategy, a 5-fold stratified cross validation with 10 repeats were applied. This classification strategy were also repeated for a one-vs-one classification of MB vs each one of other reports separately. To compare the results with an empirical chance level, a dummy classifier were also used to classify MB from other reports. This dummy classifier generated random predictions by respecting the training set class distribution.

### Neurobehavioural coupling

To evaluate the similarity between mental states’ functional connectivity patterns and the main resting state recurrent functional configurations, we extracted the five connectivity matrices preceding each probe as the functional repertoire of each specific mental state and then calculated their cosine distance to the main resting state patterns (In order to consider the effect of hemodynamic response, all the analysis were performed on the shifted versions of the connectivity matrices with time lags ranging from zero (5 matrices prior the probe) to 3 (two pre-probe and 3 post-probe matrices). Cosine distance between two sample matrices of A and B can be calculated as:

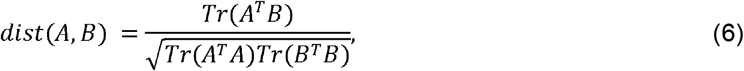

Where Tr(.) indicates trace of a matrix. Subsequently, for each mental state the distribution of distances to all four centroids were created. A generalized linear mixed effect model with gamma distribution and log link function was applied to test the relationship between the distances to each pattern and the mental states. In this model, mental state reports were considered as fixed effects and participants as random effects with sex and age as confound variables. To correct for the multiple comparison problem due to fitting the model to the distance values of different patterns separately, an effect was considered significant if its p-value was less than 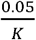, wher K is the number of patterns. In the case of a significant effect, a Tukey post-hoc test were applied to compare each pair of mental states separately.

### Global signal effect analysis

The global signal for each subject was calculated after applying the atlas and time series extraction, by averaging time series of all the ROIs. To study the effect of the global signal on the analysis results, we simply once subtracted it from the time series related to each ROI (Global Signal Subtraction (GSS)):

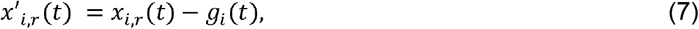

Where *i* identifies the subject, *r* identifies the ROI, and *g_i_*(*t*) is the global signal of the subject í, and once regressed it out from the ROI time series (Global Signal Regression (GSR)):

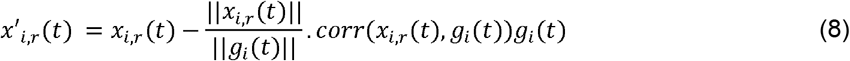

All the analysis related to the connectivity pattern extraction, their occurrence rate, and neurobehavioural coupling were also repeated in these signal versions. To study the relationship between global signal and mental states, the global signal amplitude was calculated for each mental state. The global signal amplitude was defined as the sum of the absolute value of the 5 global signal time points related to the functional repertoire of each mental state. A generalized mixed effect model with gamma distribution and inverse link function was fitted to the global signal amplitude values, considering mental states as main effect, and subjects as random effect of the model. In case of finding a significant effect, a Tukey post-hoc test was performed to compare each pair of the mental states in terms of their related global signal amplitude.

## >Data Availability

Preprocessed functional data at the level of ROI time series can be freely downloaded from: https://osf.io/3vqb6/download. The raw data are available upon request from Dr Athena Demertzi, Prof. Steve Majerus, or Dr Laurens Van Calster.

## Code Availability

All the preprocessing and analysis codes are freely available on gitlab: https://gitlab.uliege.be/S.Mortaheb/mind_blanking

## Acknowledgments

This work was supported by Belgian Fund for Scientific Research (FNRS). S.Mortaheb is a Research Fellow, A.Demertzi is a Research Associate, and S.Majerus is a Research Director at the FNRS. We also thank Dr Matthieu Koroma, Dr Camilo Miguel Signorelli, and Mr Larry D. Fort for proof reading and editing the manuscript.

